# Generative inverse design of RNA structure and function with gRNAde

**DOI:** 10.1101/2025.11.29.691298

**Authors:** Chaitanya K. Joshi, Edoardo Gianni, Samantha L. Y. Kwok, Simon V. Mathis, Pietro Liò, Philipp Holliger

**Author notes:** Equal contributions (Chaitanya K. Joshi, Edoardo Gianni, Samantha L. Y. Kwok).

## Abstract

The design of RNA molecules with bespoke three-dimensional structures and functions is a central goal in synthetic biology and biotechnology. However, progress has been limited by the challenges of designing complex tertiary interactions such as pseudoknots, as well as engineering catalytic functions—problems that have remained largely intractable for automated methods. Here we present a high-throughput generative AI pipeline for inverse design of RNA structure and function. Central to the pipeline is *gRNAde*, an RNA language model conditioned on 3D backbone structures and sequence constraints. We have validated the gRNAde pipeline in a community-wide, blinded RNA design competition on the Eterna platform, where it proved able to design complex pseudoknotted RNAs at success rates matching that of human experts (95%), while significantly outperforming other physics- and AI-based automated algorithms (70%). We further demonstrate gRNAde’s capabilities by generatively designing functional RNA polymerase ribozymes (RPR) with nearly 20% sequence divergence from the wild type RPR, discovering highly active variants at mutational distances inaccessible to rational design or adaptive walks by directed evolution. gRNAde thus provides an experimentally validated, open-source platform for automated design of complex RNA structures and accelerated engineering of complex RNA functions, providing a step towards programmable RNA catalysts and nanostructures.

Open-source code: github.com/chaitjo/geometric-rna-design

## Introduction

The unique dual capacity of RNA to both encode genetic information in its sequence and perform biochemical functions by adopting complex three-dimensional folds grants it a central role in biology^1^. RNA structure underpins a vast range of cellular processes^2^ as well as emerging biotechnologies and therapeutics^3–6^. The ability to design RNA structure and function is thus a key enabling technology for modern biology.

RNA engineering has advanced through iterative mutation and directed evolution. Recent advances in deep learning for protein design^7,8^ now open the prospect of automated RNA design. However, two fundamental and interconnected challenges need to be overcome for progress in RNA design: the design of complex three-dimensional structures and the engineering of (catalytic) functions. Here we address the first challenge by the example of RNA pseudoknots, a class of structural motifs unique to RNA, in which base-pairing of single-stranded regions with their complementary sequences creates interwoven stem-loop structures. These motifs are critical for a wide range of biological functions, from enabling viral replication in SARS-CoV-2^9^ to forming the catalytic core of ribozymes^10^, yet their topological complexity makes them notoriously difficult to predict or design computationally^11,12^. Therefore, they represent a major gap in our ability to engineer RNAs with precise three-dimensional topologies. The second, and arguably greater, challenge is to go beyond static structure and engineer complex catalytic functions. Designing active RNA enzymes (ribozymes), such as the RNA polymerase ribozymes that catalyze RNA-templated RNA copying^13–19^, is significantly more challenging than designing structure as RNA structure, although a prerequisite for function, is by itself insufficient. Finding active catalysts therefore requires navigating a vast sequence space where functional solutions are exceedingly rare. Existing computational approaches, from traditional rational design^20^ to recent deep learning models that use evolutionary information^21,22^ or base pairing constraints^23^, are often limited by handcrafted heuristics or the need for multiple sequence alignments. This leaves a critical gap for automated methods that can accurately design 3D topologies and catalytic functions directly from geometric principles, particularly for targets without evolutionary precedents.

This work introduces a comprehensive solution to these twin challenges. We describe a new, generative AI pipeline, called *gRNAde*^1^, that learns the principles of RNA folding directly from 3D structural data, and is integrated into a complete design-build-test framework. The foundational architecture, which leverages an RNA language model to generate sequences conditioned on 3D backbone structures has been described previously^24,25^, but its practical ability to solve major experimental challenges in RNA engineering had remained untested. Here we describe a systematic validation of gRNAde’s performance in both a blinded structural design competition for pseudoknot structures on Eterna^26^ as well as a complex functional design campaign for novel RNA polymerase ribozyme variants, demonstrating its superior performance as a generalizable, automated, and open-source platform for RNA design.

## Results

### A generalizable pipeline for high-throughput RNA design

To address the challenges of RNA design at scale, an automated pipeline was developed to integrate the generative capabilities of gRNAde^24,25^ with high-throughput computational screening (**Figure 1**). This workflow provides a practical and efficient method for identifying promising RNA sequences for experimental validation.

**Figure 1.**
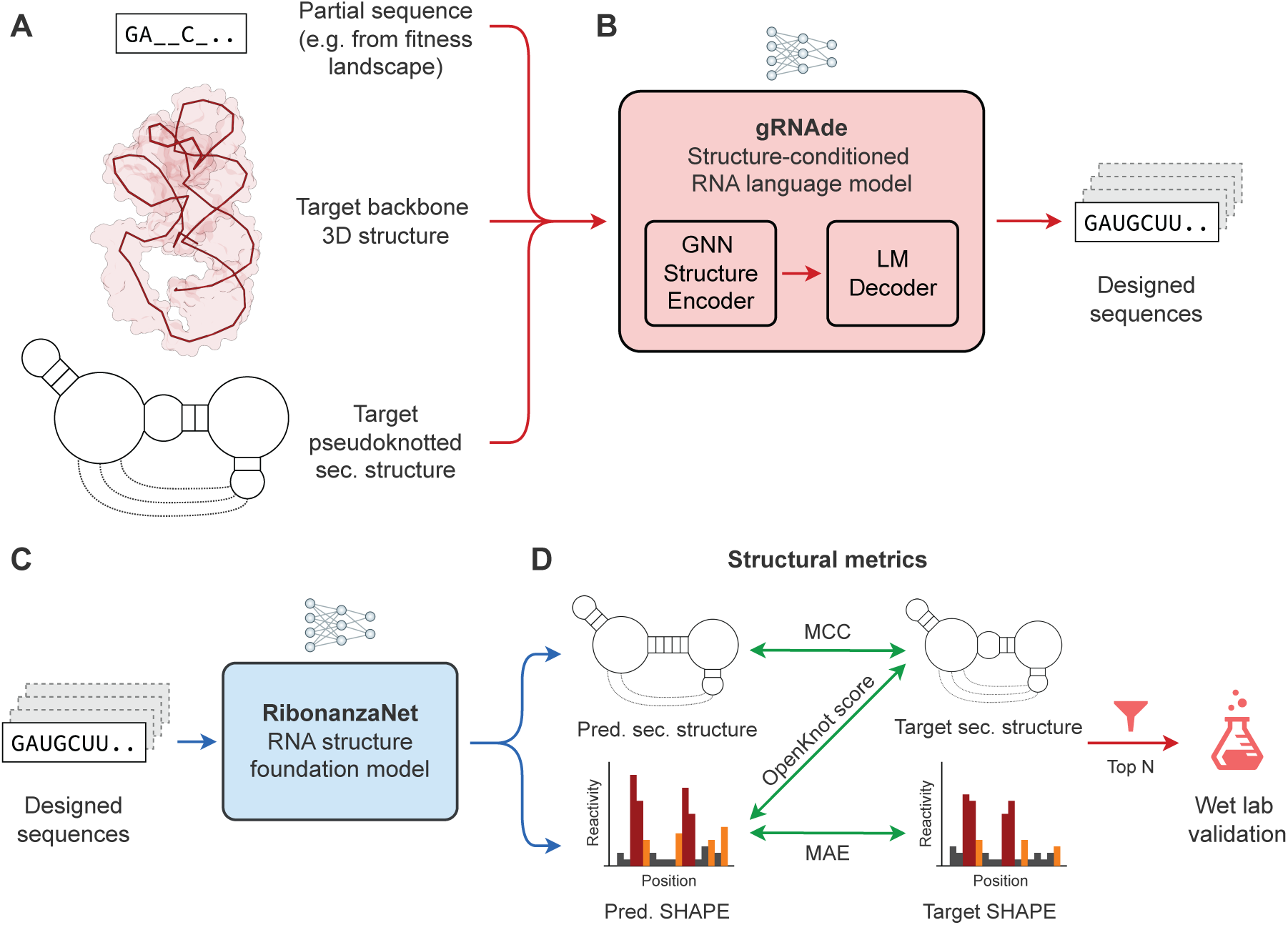
The gRNAde high-throughput pipeline for generative RNA design. The automated workflow integrates deep learning-based sequence generation with computational screening to identify optimal candidates for experimental validation. **A.** The pipeline takes multi-modal design constraints as input, optionally including a target pseudoknotted secondary structure, a target 3D backbone structure, and partial sequence constraints such as those derived from fitness landscapes. **B.** gRNAde, a structure-conditioned RNA language model, uses these constraints to generate a large and diverse library of candidate sequences, typically on the order of one million. These candidates are then passed to the computational filtering stage. **C.** In the filtering stage, each designed sequence is evaluated by RibonanzaNet, an RNA structure foundation model. RibonanzaNet predicts the secondary structure and a per-nucleotide SHAPE chemical reactivity profile for each candidate. **D.** RibonanzaNet predictions for each design are scored against the target secondary structure and SHAPE profile using metrics such as the Matthews Correlation Coefficient (MCC), Mean Absolute Error (MAE), and the OpenKnot Score. The top-ranked designs are then selected for wet-lab synthesis and validation.

Conceptually, gRNAde is a generative language model for RNA, translating a multi-modal structural ‘prompt’ into novel sequences predicted to adopt a desired fold. This design specification, analogous to a textual prompt for a large language model, is highly flexible; it can consist of a target pseudoknotted secondary structure, 3D backbone coordinates, and partial sequence constraints that must be preserved (**Figure 1A**). Unlike traditional physics-based or heuristic methods, gRNAde is a data-driven model whose design principles are learned directly from the thousands of experimentally determined RNA structures in the Protein Data Bank^28^. Its core advance is the direct integration of 3D geometric information using Graph Neural Networks (**Figure 1B**), allowing it to capture the subtle tertiary interactions and non-canonical pairings that define complex RNA folds—features inaccessible to methods restricted to secondary structure representations.

The full design pipeline proceeds in a three-step workflow. First, in the generation stage, gRNAde is used to produce a large and diverse library of candidate sequences, typically on the order of one million, for a given structural target. Second, in the screening stage, each generated sequence is evaluated using RibonanzaNet^27^, an RNA structure foundation model. RibonanzaNet serves as a high-throughput computational proxy for experimental characterization by predicting per-nucleotide chemical reactivity profiles and pseudoknotted secondary structure (**Figure 1C**). These reactivity profiles, which measure the accessibility of each nucleotide to chemical modification, are highly sensitive reporters of an RNA’s folded 3D structure and flexibility^29,30^. The choice of RibonanzaNet is motivated by its training on a massive and diverse dataset of both natural and synthetic RNAs, making it well-suited to evaluate novel, designed sequences. Third, in the selection stage, the computationally screened candidates are scored and ranked (**Figure 1D**). Key metrics include the Matthews Correlation Coefficient (MCC) between the predicted secondary structure and the target, and the Mean Absolute Error (MAE) between the predicted chemical reactivity profile and a target profile. This filtering process distils the massive initial library down to a small, high-quality set of candidates for synthesis and wet-lab validation. This entire pipeline is highly efficient, capable of generating and screening one million designs in under 12 hours on a single GPU, making it a powerful tool for exploring RNA sequence space on an unprecedented scale.

### Expert-level design of pseudoknotted RNA structures

To rigorously assess the pipeline’s ability to solve the “RNA Structure Challenge”, gRNAde was entered into the Eterna OpenKnot Benchmark, a community-wide, blinded competition to design sequences for complex RNA pseudoknot targets. The benchmark is exceptionally challenging, with targets spanning a wide range of biologically critical structured RNA classes, including riboswitches, viral elements, ribozymes, and synthetic structures, ensuring a robust test of the method’s generalizability. Submitted designs are experimentally synthesized and evaluated at Stanford University, with performance quantified by an “OpenKnot Score” derived from high-throughput chemical probing data; a score above 90 indicates a high-confidence successful design. This benchmark provides an unbiased comparison against both state-of-the-art automated methods and the collective intelligence of human experts^26^.

Across 20 challenging targets of up to 100 nucleotides (nts) in length (OpenKnot Round 3), the fully automated gRNAde pipeline achieved a 100% success rate on natural RNAs and a 90% success rate on synthetic targets, on par with expert human designers who also achieved 100% and 90% success rates, respectively (**Figure 2A-D**). In stark contrast, the state-of-the-art physics-based method, Rosetta^7^, only succeeded on 40% of natural and 70% of synthetic targets. RFdiffusion^31^, the next-best public AI method, also struggled with success rates of 80% and 60%, respectively. These difficulties were particularly pronounced with longer targets of 240 nts in OpenKnot Round 4 (**Supplementary Figure 1**). Here, gRNAde remained competitive with experts, solving 67% of natural and 70% of synthetic targets, while Rosetta was not scalable enough to be evaluated and RFdiffusion succeeded on only 33% and 50%, respectively. Full results of the competition, including additional compensatory mutagenesis and cryo-EM experiments, will be detailed in a forthcoming joint publication and signify a key milestone in biomolecular design: our fully automated gRNAde pipeline has matched human experts across a blinded, challenging set of RNA structure design tasks.

**Figure 2.**
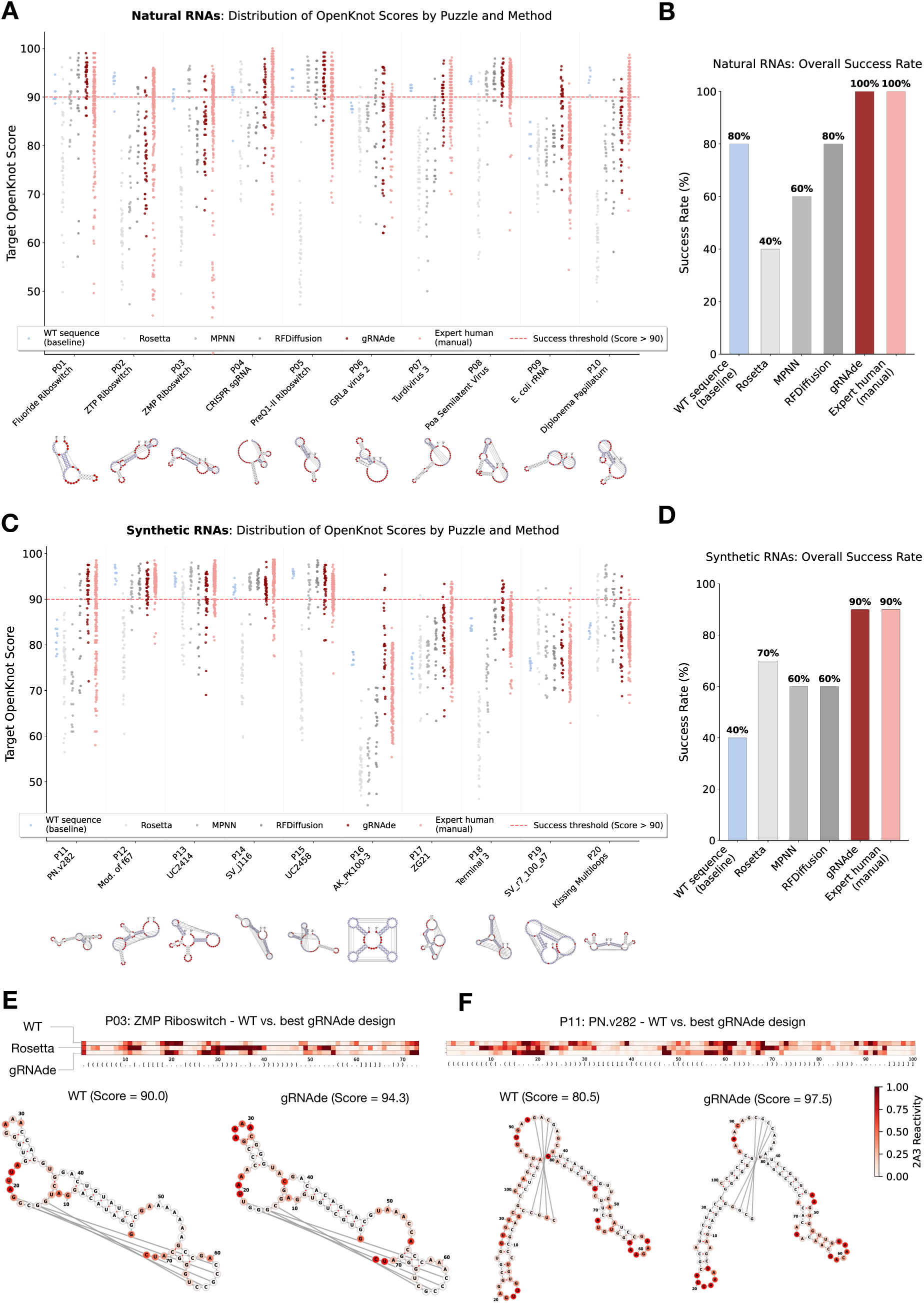
gRNAde achieves expert-level accuracy in the Eterna OpenKnot Benchmark for RNA pseudoknot design. Performance of wildtype sequences, Rosetta^12^, MPNN^32^, RFdiffusion^31^, gRNAde, and expert human designers in the Eterna OpenKnot Round 3 challenge, which targeted pseudoknotted RNAs of up to 100 nucleotides. **A, B.** Results for 10 natural RNA targets. **C, D.** Results for 10 synthetic RNA targets. Left-sided panels A and C show the distribution of OpenKnot scores for individual designs across all puzzles (success threshold > 90, red dashed line). Right-sided panels B and D show the overall success rate, defined as the percentage of puzzles for which at least one design scored above 90. gRNAde achieves success rates of 100% (natural) and 90% (synthetic), matching expert human performance and substantially outperforming both Rosetta (physics-based) and RFdiffusion (next best AI method). **E, F.** Molecular validation of design success through chemical probing. Nucleotides are overlaid on the target secondary structures and colored by reactivity, with darker reds indicating higher reactivity and greater accessibility for unpaired positions. Conversely, nucleotides part of base pairs and pseudoknots are expected to have lower reactivity. **E.** For the natural ZMP Riboswitch target, the best gRNAde design (right, score = 94.3) shows a chemical reactivity profile largely consistent with the target fold, whereas the wildtype sequence (left, score = 90.0) shows anomalous reactivity in the loop region around position 50-55, suggesting misfolding. **F.** For the synthetic PN.v282 target, the gRNAde design (right, score = 97.5) again shows a superior reactivity pattern compared to the wildtype sequence (left, score = 80.5), which exhibits high reactivity in various paired regions, indicating disrupted base pairing.

Notably, gRNAde’s designs consistently outperformed the wildtype sequences from which the natural targets were derived. Wildtype sequences achieved success rates of only 80% on short targets and a striking 0% on long natural targets. This finding underscores an important principle: while natural sequences may form alternative conformations, gRNAde’s designs are often more “idealized” solutions for a target structure. The molecular basis for this superior performance is evident in the experimental data. For the natural ZMP Riboswitch target, the chemical reactivity profile of the gRNAde design precisely matches the expected pattern for the target fold, while the wildtype sequence shows anomalous reactivity patterns suggesting an alternative conformation (**Figure 2E**). This trend holds for synthetic targets as well; for the PN.v282 puzzle, the gRNAde design again shows a near-perfect reactivity profile, whereas the wildtype exhibits high reactivity in paired regions, indicating a failure to form the correct idealized structure (**Figure 2F**). This direct experimental evidence confirms that gRNAde can generate complex target topologies, whereas the tested natural pseudoknot sequences display topologies deviating from the idealized target due to multiple evolutionary selective pressures.

### Generative design of functional RNA enzymes

Having demonstrated expert-level performance at designing complex structural topologies, the next challenge was to address the design of function. The gRNAde pipeline was applied to engineer the 5TU catalytic subunit of a triplet-based RNA polymerase ribozyme^33^, a highly complex ribozyme that serves as a model for RNA self-replication in the context of origins of life research. This design task is demanding as it requires preserving a delicate network of tertiary contacts, including two essential kissing-loop interactions that preorganize the enzyme’s active site (**Figure 3A, B**), a challenge that goes far beyond simple secondary structure conservation. The scientific goal was not merely to maintain and if possible improve enzyme function, but to test the limits of its functional sequence diversity^22,34,35^, i.e. the maximal edit distance of the 5TU “quasispecies” in RNA sequence. This has direct implications for the plausibility of RNA-based self-synthesis and the emergence of early life^36,37^. This was achieved by using gRNAde to perform large-scale generative “jumps” in sequence space (**Figure 3C**), aiming to discover functional variants at mutational distances far beyond those accessible through adaptive walks using conventional directed evolution or simple rational design. By traversing vast sequence distances in a single step, gRNAde shortens the time investment of multiple selection rounds and accesses variants that comprise compensating negative mutations that are challenging to access by evolution. This has the potential to accelerate or offer novel starting points for directed evolution.

**Figure 3.**
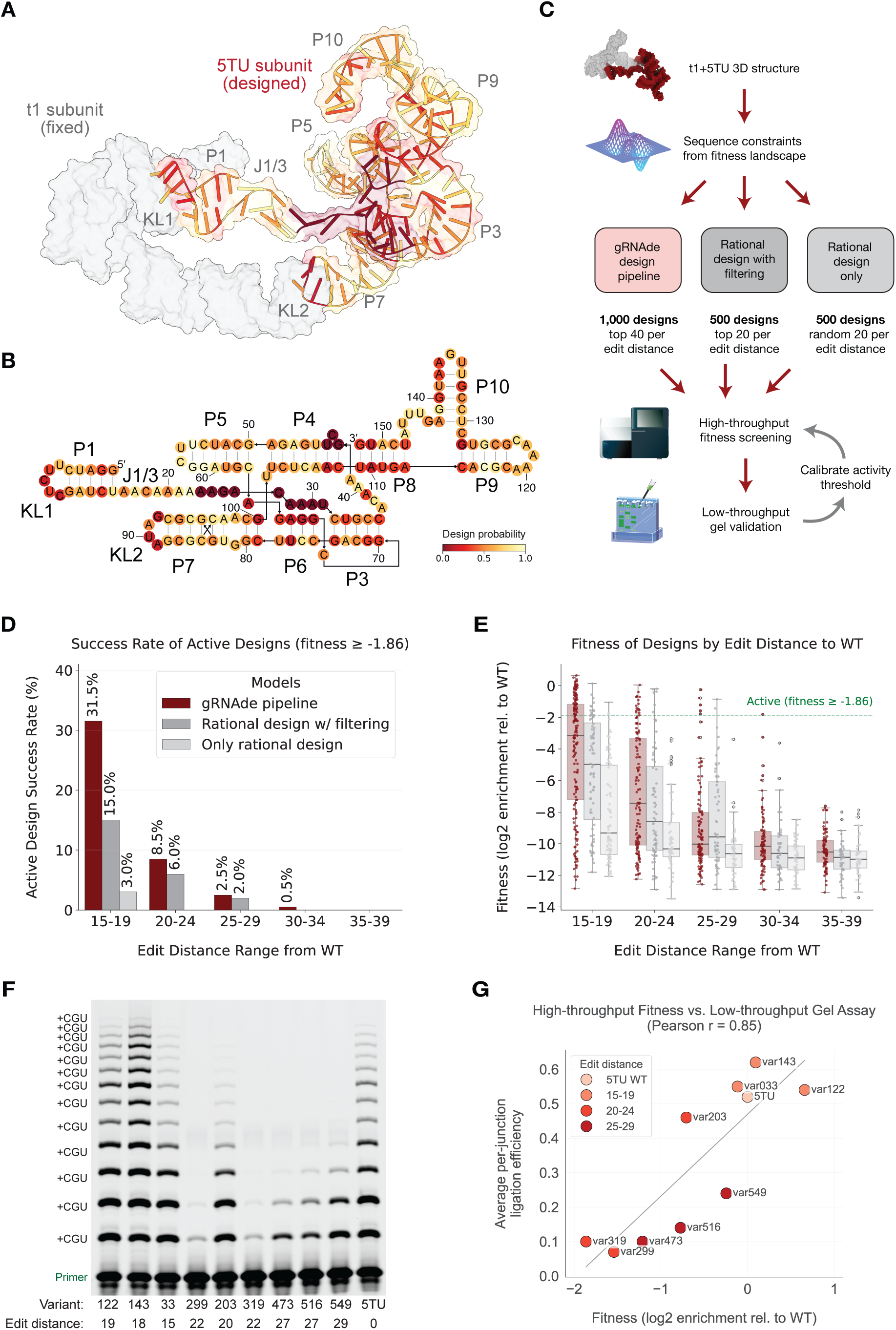
Generative design and functional validation of RNA polymerase ribozymes. The gRNAde pipeline was used to design functional variants of the triplet-based RNA polymerase ribozyme (TPR), substantially outperforming rational design baselines. **A.** Cryo-EM structure of the TPR heterodimer (PDB: 8T2P), showing the catalytic 5TU subunit (colored), which was the target for generative design, and the auxiliary t1 subunit (grey), which was held constant. Position-specific design probabilities, derived from experimental fitness landscape data^33^, are mapped onto the 5TU structure. Lighter yellows indicate regions with a high probability of being re-designed by gRNAde, while critical functional sites were constrained to the wildtype sequence (indicated in darker reds). **B.** Position-specific design probabilities mapped onto the 5TU secondary structure. **C.** Workflow for design and validation of 5TU variants. The 3D backbone structure, along with constraints sampled from the fitness landscape data, were input to the full gRNAde pipeline (Figure 1) as well as two baselines: rational design with the same computational filtering as gRNAde, and rational design without filtering. A library of 2,000 total designs was screened via a high-throughput functional assay. The native 5TU and 9 gRNAde designs were further validated using a low-throughput gel, which was then used to calibrate the activity threshold for the high-throughput data. **D.** Success rate of generating active designs (fitness ≥ -1.86, corresponding to variant 319) binned by mutational distance from the wildtype sequence. At 15–19 mutations, the gRNAde pipeline achieves a 31.5% success rate, substantially outperforming filtered rational design (15.0%) and unfiltered rational design (3.0%). **E.** Fitness distributions for all functional designs across mutational distances. The fitness of gRNAde designs is consistently higher than that of the baseline methods, with many variants exceeding the activity threshold (fitness ≥ -1.86). **F.** Low-throughput primer extension reactions on a template encoding 14 repeats of CGU, using the top 9 gRNAde-designed variants by fitness in the high-throughput assay. Variant identity and edit distance from wildtype 5TU is labelled below the gel. The gel confirms the activity of all gRNAde variants, with variants 122, 143, 33, and 203 showing activity comparable or better than the native 5TU ribozyme. **G.** Correlation between high-throughput functional assay and low-throughput gel for the native 5TU sequence and 9 gRNAde-designed variants. The fitness scores are highly correlated with the average per-junction ligation efficiency from the gel (Pearson r = 0.85), and the fitness of the least active variant 319 is used as an activity threshold for the high-throughput assay, as it demonstrates some ligation activity on the gel.

The gRNAde pipeline dramatically outperformed baseline methods in discovering active ribozymes. For designs with 15-19 mutations relative to the 152 nt wildtype sequence, gRNAde yielded a 31.5% success rate in identifying active variants in a high-throughput sequencing-based functional assay (**Figure 3D**). In contrast, a “rational design” baseline that respects secondary structure but lacks 3D awareness, even when coupled with the same RibonanzaNet filtering used in gRNAde, achieved only 15% success rate. This is a more than two-fold improvement and demonstrates that the performance gain stems directly from gRNAde’s superior generative capability, not just the downstream filtering. Furthermore, without filtering, gRNAde showed a more than ten-fold improvement over rational design, which had a success rate of just 3%.

Furthermore, gRNAde discovered not only more functional variants but also variants with higher catalytic activity on model templates. The fitness distribution of active gRNAde designs was generally superior to that of the rational design baselines (**Figure 3E**). Three active gRNAde variants each for mutational distance ranges 15-19, 20-24, and 25-29 were further validated via a low-throughput gel assay, showing high Pearson correlation coefficient of 0.85 with the high-throughput fitness (**Figure 3F, G**). Notably, variants 122 and 143 differed from the wildtype by 18 and 19 mutations, yet exhibited an 1.6-fold and 1.1-fold enrichment in the high throughput assay, respectively. gRNAde retained activity even with up to 28 mutations (**Supplementary Figure 3**). These results showcase the combination of sequence novelty and improved functionality achieved with gRNAde.

### Mechanistic insights from gRNAde’s generative design strategy

To understand the basis for gRNAde’s superior performance across diverse RNA design tasks, we analyzed the mutational patterns in active ribozyme designs from the gRNAde pipeline compared to rational design with filtering. Rational design tended to mainly mutate canonical base-paired positions while conserving unpaired loops, a strategy that mainly preserves secondary structure but fails to account for essential tertiary interactions (**Figure 4B**). gRNAde, in contrast, generated active designs with a more balanced mutational profile (**Figure 4A**), frequently altering nucleotides in unpaired regions and loops, altering nucleotides in four unpaired regions: J1/3 (single-stranded template-binding interface positioned by kissing loops), the loop region of P5 (structural scaffold of catalytic core), the loop region of P9 (structural scaffold of the extension domain), and the loop region of P10 (makes critical substrate contacts^16^ and shows dynamic movement towards active site) (**Figure 4C-E**). gRNAde can successfully mutate these four regions with diverse functions, demonstrating that by training on a diverse corpus of 3D structures, it has learned sophisticated, non-local structure-function relationships that go far beyond simple base-pairing rules, allowing it to successfully navigate a complex functional landscape.

**Figure 4.**
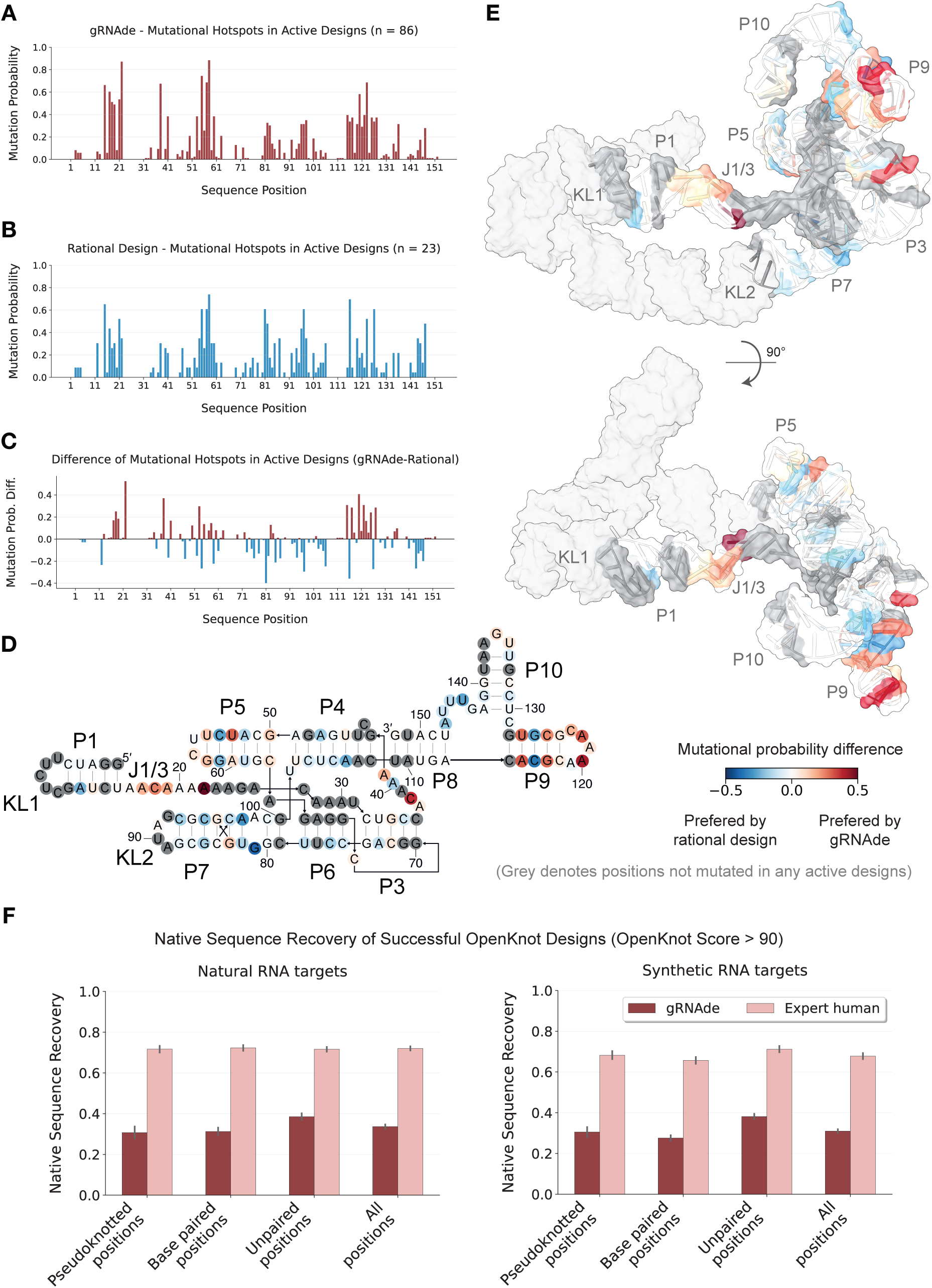
Mechanistic analysis of gRNAde design strategies. The analysis compares gRNAde’s design strategy against rational design and human experts, demonstrating its capacity to learn non-local, 3D-informed structure-function relationships and achieve highly sequence-divergent yet structurally accurate designs. **A-C**. Per-position mutation probability (“hotspots”) for active RNA polymerase ribozyme designs generated by gRNAde (A) and rational design with filtering (B), and the difference between them (C). **D, E**. The difference in mutation probability is mapped onto the secondary structure (D) and tertiary structure (E) of the catalytic subunit 5TU that was the target of design. This reveals distinct design strategies: Rational design preferentially mutates canonical base-paired positions (blue) in active designs, whereas gRNAde identifies novel hotspots in structurally complex, unpaired regions (red), particularly near the template-binding site (J1/3) as well as ends of helices P5, P9, and P10. **F.** Median native sequence recovery of successful designs from gRNAde and expert human designers in OpenKnot Round 7, presented by position type (pseudoknotted, paired, unpaired, and all positions) for natural (left) and synthetic (right) RNA targets. Across all position types, gRNAde designs exhibit significantly lower median native sequence recovery (32%) compared to human experts (72%). This demonstrates gRNAde’s capacity to achieve large generative jumps in sequence space while matching expert accuracy at forming the target structure.

To further contextualize gRNAde’s design strategy against human expertise, we revisited the OpenKnot Benchmark to analyze successful designs in Round 7 by computing their native sequence recovery from the wildtype or starting sequence (**Figure 4F**). The analysis revealed that while gRNAde matched the structural accuracy of human experts, its designs were significantly more distant in sequence space. The median native sequence recovery for gRNAde designs was 32%, substantially lower than the 72% observed for human experts. This divergence demonstrates that unlike human designers, who exhibit a strong bias toward smaller more conservative edits close to the native sequence, gRNAde can successfully perform generative jumps in sequence space for diverse RNA targets. This capability underscores gRNAde’s key advantages: its data-driven understanding of structurally complex, unpaired regions and its capacity to retain the target structure across vast sequence distances.

## Discussion

This work has introduced and experimentally validated gRNAde, a general-purpose inverse design pipeline for RNA. gRNAde provides solutions for two of the most significant and long-standing challenges in the field. First, in a blinded, community-wide competition, the automated pipeline successfully designed complex RNA pseudoknot structures with an accuracy matching that of human experts, establishing a new state-of-the-art for structural RNA design. Second, the pipeline was used to generatively explore the functional landscape of a complex RNA polymerase ribozyme, discovering highly active ribozymes at large sequence distances from any known functional variant. This dual success on fundamentally different problems—one focused on folding and structural accuracy, the other on function, embodying both structure and dynamics—validates the power and generality of the approach.

These findings have broader implications for our understanding of both natural and engineered biological systems. The OpenKnot results, where gRNAde’s designs proved more stable for targeted structural goals than their native counterparts, suggest that data-driven optimization can uncover solutions that are more idealized than those found in biology, where evolution operates under a multitude of competing constraints. Similarly, the discovery of a diverse and functional ribozyme quasispecies, with active variants differing by nearly 20% of their sequence, demonstrates that the functional sequence space for complex RNA is likely larger than anticipated comprising (presumably) structurally-similar variants at mutational distances that would be challenging to access by directed evolution. Generative models like gRNAde provide a powerful new tool to explore this vast, uncharted territory, providing an alternative to local exploration by directed evolution or the limitations of human-centric rational design.

Beyond its direct applications, this work highlights the potential for a virtuous cycle in computational biology, where the success of gRNAde in generating vast libraries of high-quality designs has enabled the creation of new, large-scale datasets. For example, a follow-on collaboration used gRNAde to generate 68 million plausible sequences for 1.6 million pseudoknotted structures, a dataset orders of magnitude larger than previously possible. This dataset then trained RibonanzaNet 2 with significantly improved accuracy^38^, creating a powerful feedback loop where generative models produce data to train better structure prediction models, which can then be incorporated back into the design pipeline as more accurate filters, further accelerating progress.

Looking forward, as advances in experimental techniques like cryo-EM continue to expand the universe of known RNA structures^39,40^, the data foundations for RNA design models will grow, making pipelines like gRNAde increasingly powerful and bringing the goal of programmable RNA biology closer to reality. We anticipate that gRNAde will be widely useful, enabling the engineering and design of novel riboswitches, ribozymes, and RNA nanostructures. To accelerate progress and enable broader community use, gRNAde is fully open-source at github.com/chaitjo/geometric-rna-design.

## Methods

### gRNAde model and training

The gRNAde model is a structure-conditioned autoregressive language model for RNA sequence design. We have previously described the model architecture^24^ and workflow^25^ in detail. Here, we made key modifications to the input representation and training procedure to enable design from both secondary and tertiary structure, as well as sequence constraints.

The model takes as input a target RNA’s pseudoknotted secondary structure, a 3D backbone structure, and partial sequence constraints. Each of these three input modalities is optional and the model can flexibly handle missing inputs. Each nucleotide is represented as a node in a geometric graph, featurized using a 3-bead coarse-grained representation (P, C4’, and N1/N9 atoms)^41^ and local geometric properties^42^. A key architectural modification from the initial version of gRNAde is the explicit integration of secondary structure information into the graph. Each node is connected to its neighbors by edges of three types: (a) its 32 nearest neighbors along the RNA backbone (1D); (b) all nodes involved in base pairs and pseudoknots from the secondary structure based on DSSR^43^ (2D); and (c) its 32 nearest neighbors in 3D space, if a 3D structure is provided. Edge features encode the edge type, relative distances and orientations between nodes. The core of the model is an SE(3)-equivariant Graph Neural Network^44^ (GNN) encoder composed of Geometric Vector Perceptron (GVP-GNN) layers^45^, which processes the geometric graph to produce a latent representation for each nucleotide. An autoregressive GVP-GNN decoder then uses these representations to predict the nucleotide sequence from 5’ to 3’. We use 4 layers each for the encoder and decoder. If a partial sequence is provided as a constraint, the decoder fixes the identity of those nucleotides via logit biasing and designs the remainder of the sequence, similar to ProteinMPNN^7^.

The model was trained on a dataset of 4,211 RNA structures with up to 500 nucleotides, covering a diverse set of topologies from the Protein Data Bank^46^ at resolution <= 4.0 A, curated by RNASolo^28^ with a temporal cutoff of 31 October 2023. To enable design without complete 3D information, we introduced a novel training strategy: for a randomly selected 50% of training examples at each training epoch, the 3D backbone coordinates and corresponding 3D edges were dropped out. This forces the model to learn the implicit 3D structural information contained within the 1D and 2D graph connectivity alone. As a result, the trained model can generate high-quality designs even when a 3D structure is unavailable or unreliable. This capability was critical for designing the synthetic targets in the OpenKnot Benchmark, where only a target secondary structure was available.

### Computational design and screening pipeline

For each design task, an initial library of approximately 1 million candidate sequences was generated using gRNAde by sampling from the model’s output distribution with varying temperatures (0.1 to 1.0) and random seeds. This allows the model to balance high-confidence, lower-diversity sampling (low temperature) with broader exploration of the solution space (high temperature).

Each generated sequence was then screened using RibonanzaNet, an RNA structure foundation model^27^. We chose RibonanzaNet as its predictions of per-nucleotide chemical reactivity and secondary structure serve as a more reliable computational proxy for experimental folding than current 3D structure predictors. While we initially evaluated 3D self-consistency using RhoFold^47^, this metric was excluded after control experiments revealed that the predictor failed to recover the correct backbone geometry for native sequences in the Eterna OpenKnot Benchmark. Given RhoFold’s inability to reliably fold the ‘ground truth’ inputs, we concluded that current 3D prediction tools^48–50^, which perform equivalently to RhoFold^51,52^, may not be sufficiently robust to serve as filters for de novo designs.

The screening process involved predicting a per-nucleotide chemical reactivity profile (for the 2A3 modifier) and a secondary structure for each sequence. Designs were then ranked based on two primary scores: (1) a Secondary Structure Score, calculated as the Matthews Correlation Coefficient (MCC) between the predicted and target secondary structures. For natural targets, we applied a stringent filter, retaining only sequences with an MCC > 0.9 to ensure a high likelihood of correct folding; this criterion was omitted for synthetic targets as RibonanzaNet’s secondary structure predictor was fine-tuned only on natural sequences. (2) a Chemical Reactivity Score, calculated as the Mean Absolute Error (MAE) between the predicted reactivity profile and a target profile (either from experimental data or predicted from the native sequence). For the OpenKnot Benchmark, instead of the MAE, we used an *in silico* OpenKnot Score between the predicted reactivity profile and the target secondary structure.

The entire pipeline is highly efficient; generating and scoring one million sequences for a design campaign takes under 12 hours on a single NVIDIA A100 GPU. After removing duplicates, this process typically yields hundreds of thousands of unique sequences, from which the top-ranked candidates were selected for experimental validation. This high-throughput capability contrasts sharply with physics-based methods like Rosetta^12^, where generating a single design can take over 24 hours on a CPU, making large-scale exploration computationally prohibitive.

### Eterna OpenKnot Benchmark blinded competition

The gRNAde pipeline was benchmarked in the Eterna OpenKnot Benchmark rounds 7a and 7b, targeting 40 diverse pseudoknotted RNA structures. We submitted two distinct sets of 20 designs per target—one conditioned on both the target 3D backbone and pseudoknotted secondary structure, and one conditioned solely on the secondary structure—to assess the necessity of explicit 3D inputs. For synthetic targets lacking experimental structures, predicted backbones provided by the Eterna organizers were utilized for the 3D-conditioned set.

Experimental validation was performed independently by the Eterna organizers at Stanford University. Synthesized RNAs were subjected to high-throughput chemical probing using SHAPE (Selective 2’-Hydroxyl Acylation analyzed by Primer Extension) with the reagent as 2-aminopyridine-3-carboxylic acid imidazolide (2A3)^53,54^. The resulting reactivity data was used to calculate an OpenKnot Score^27^ (between 0-100), which quantifies the agreement between the experimental SHAPE profile and the desired profile of unpaired and paired nucleotides for the target structure. An OpenKnot Score greater than 90 was considered a high-confidence successful design that is likely to fold into the target structure.

To provide a direct automated baseline for comparison, the organizers also independently submitted 20 designs using Rosetta’s RNA inverse design protocol^12^, the current state-of-the-art physics-based method for 3D RNA design. As an additional sanity check, the organizers evaluated 10 replicates of the wildtype (native) RNA sequence for each puzzle. While wildtype sequences are expected to achieve high OpenKnot scores, particularly for natural RNAs, they may not represent the optimal sequence for structure formation, which is precisely what the design challenge aims to discover. The competition also included other, contemporaneous AI-based methods from Stanford University^55^, University of Washington^31,32^, and Texas A&M University (not public). A full head-to-head comparison of all automated methods will be presented in a forthcoming publication.

### RNA polymerase ribozyme design

Functional variants of the 5TU catalytic subunit of the triplet polymerase ribozyme^16^ were designed based on its cryo-EM structure^33^, while the sequence of the t1 auxiliary subunit was held constant. To guide the design process, we defined position-specific mutation probabilities for the 5TU subunit derived from the fitness landscape data of McRae et al. This procedure combined two complementary metrics:

1. Maximum single-mutant fitness, showing tolerance to point mutations. Fitness values were uniformly binned into 3 categories (1 = low, 2 = medium, 3 = high).
2. Higher-order mutant combinability score^56^, quantifying how well mutations at each position can be combined with other mutations to create active variants. Combinability values were uniformly binned into 3 categories (1 = low, 2 = medium, 3 = high).

To yield the final design probability for each position (**Figure 3B**), the binned single-mutant fitness and combinability scores were summed and divided by 6. This yields a probability value between 0.0 to 1.0 for each position. To preserve essential catalytic activity, key functional residues at the catalytic site (positions 41-43), template binding region (positions 22-24), and triple helix-forming adenosines (positions 25-30) were constrained to the wildtype identity by setting their design probability to zero.

The design probabilities are used as stochastic constraints for positions to be mutated when creating a batch of designs with gRNAde. 1 Million designs were generated with gRNAde and filtered to select 1,000 candidates spanning mutational distances of 15 to 40 from the wildtype sequence (40 designs per mutational distance). This represents a significant extension beyond the coverage of the fitness landscape data from McRae et al., which had almost negligible sampling beyond 6 mutations.

To isolate the contributions of gRNAde’s sequence generation versus our computational filtering pipeline, we implemented two rational design heuristics as baselines. The first, **‘rational design only’**, involved generating 1 million sequences using position-specific nucleotide identity sampling that respected base-pairing constraints, from which 500 designs were selected randomly (20 per mutational distance). The nucleotide identity for unpaired positions is sampled randomly. The second, **‘rational design with filtering’**, applied our full RibonanzaNet computational filtering pipeline to the same pool of 1 million rationally generated sequences to select the top 500 candidates. This experimental design allows for a direct comparison of gRNAde against commonly used rational approaches while ablating the specific contribution of the generative model.

### RNA polymerase ribozyme activity assays

The designed 5TU variants from gRNAde, rational design with filtering, and rational design only were synthesized as a pooled DNA oligonucleotide library (Twist Biosciences, termed gRNAde_library henceforth) of 1989 sequences. This accounts for duplicates and additionally includes the native 5TU sequence. The library was subjected to an *in vitro* activity selection assay that measures templated RNA synthesis.

### Preparation of DNA templates for transcription

PCR amplification of gRNAde_library was performed using Platinum SuperFi II PCR Master Mix (Invitrogen). Each reaction had a total volume of 50 μL and included 1 μM of primers forceGG and HDVrec along with gRNAde_library that was used as a template. The thermal cycling protocol involved an initial denaturation at 98 ℃ for 30 seconds, followed by 10 cycles of: denaturation at 98 ℃ for 10 seconds, primer annealing at 55.8 ℃ for 10 seconds, and extension at 72 ℃ for 15 seconds. This was followed by a final extension at 72 ℃ for 5 minutes, then held at 10 ℃. 4 μL of the PCR mixture was then run on a 4% UltraPure agarose (Life Technologies Ltd) gel to check that the size of the amplified construct is correct. The resultant amplified DNA library was then purified using a PCR purification kit (Qiagen).

To add on the T7 promoter sequence, the flexible linker, and the templates for copying, the purified DNA library was then used as template in another round of PCR. This was also conducted using Platinum SuperFi II PCR Master Mix (Invitrogen) under similar conditions using the primers HDVrt and 5T7HP6×6AUA_forceGG. The 5T7HP6×6AUA_forceGG primer allows for the addition of the T7 promoter sequence, the template encoding 6 AUA triplets to be copied by the ribozyme, and the flexible linker made of 6x(AACA) repeats. The HDVrt primer allows for the introduction of a HDV ribozyme 3′-end cassette. To vary the linker length and the template used to measure the ribozyme activity, this was repeated with 5T7HP12×6AUA_forceGG (template encodes 6 AUA triplets, linker contains 12x(AACA) repeats), 5T7HP6×6GAA_forceGG (template encodes 6 GAA triplets, linker contains 6x(AACA) repeats), and 5T7HP12×6GAA_forceGG (template encodes 6 GAA triplets, linker contains 12x(AACA) repeats) replacing 5T7HP6×6AUA_forceGG. The sizes of the constructs were also checked on a 4% UltraPure agarose (Life Technologies Ltd) gel and the DNA constructs were then purified using a PCR purification kit (Qiagen).

### Preparation of RNA constructs

The four DNA templates were then transcribed following an RNA transcription protocol based on the Megascript method^57^ using T7 RNA polymerase. The 50 μL reaction mixture contained 40 mM Tris buffer at pH 8, along with 10 mM DTT, 2 mM spermidine, and 20 mM MgCl. Each nucleotide triphosphate was added at 7.5 mM concentration. The DNA template was included at approximately 5 pmoles. The reaction also included 0.01 units/μL yeast inorganic pyrophosphatase (Thermo Fischer Scientific) and approximately 72 μg/mL T7 RNA polymerase (expressed and purified in house). After incubating overnight at 37 ℃ (12-16 hours), the mixture was treated with Turbo DNase (Thermo Fischer Scientific) at 0.1 units/μL for one hour.

To increase the efficiency of HDV cleavage, the *in vitro* transcription reactions were then subjected to temperature cycling. Transcription reactions were first incubated at 50 ℃ for 1 minute and then placed on ice for 3 minutes. Subsequently, for every 50 μL transcription reaction, 20 μL of gel loading buffer (95% formamide, 25 mM EDTA, bromophenol blue) was added, and the mixture was incubated at 50 ℃ for another minute. After placing the mixture on ice for 3 minutes, 120 μL of gel loading buffer was added, and the mixture was heated to 94 ℃ for 5 minutes. Then, the samples were loaded into a PAGE gel for purification.

After heat denaturation at 94 ℃ for 5 minutes, samples were loaded onto a denaturing polyacrylamide gel made of a 10% acrylamide/bis-acrylamide 19:1 solution (Severn Biotech), 8M urea, 1x Tris-borate-EDTA buffer (89 mM Tris pH 7.6, 89 mM boric acid, 2 mM EDTA). Electrophoresis was performed at 30W constant power using a Cambridge Electrophoresis EV400 system using bromophenol blue as a marker for migration. Target bands were visualized by UV shadowing and cut out from the gel. The excised gel pieces were crushed using a 1000 μL pipette tip and soaked in Tris-EDTA buffer (10mM Tris pH 7.4, 1mM EDTA) to create a suspension. Following one freeze-thaw cycle (dry ice freezing, 50 ℃ thawing) and rotation in the cold room overnight, the solution was filtered through 0.22 μm filters (Costar), and the RNA was extracted using ethanol precipitation with 73% ethanol. Final product concentration was measured using a Nanodrop ND-1000 spectrophotometer (Thermo Fisher Scientific) at 260nm using sequence-specific extinction coefficients calculated with the Oligocalc software^58^.

### High-throughput screening of polymerase ribozyme activity

To test the activity of the designed polymerase ribozyme variants a single round of *in vitro* selection for primer extension was carried out on pooled libraries of ribozyme sequences. The library composition before and after a single selection round was assessed via next-generation sequencing, and relative enrichment values were computed as proxies for fitness.

A single selection round was carried out similarly to previous reports^16,33^, where ribozyme variants are tethered via a flexible RNA linker to a template. Completion of the template copying leads to covalent linkage of the synthesized product to the ribozyme responsible to the synthesis, ensuring genotype-phenotype linkage. Active variants can therefore be enriched by selective pull-down and denaturing PAGE purification of the full-length product-ribozyme conjugates.

The selection round reactions were set up as follows: 5 pmoles of the 6×6AUA library (contains both the 5TU variants to be tested and the 6 AUA template) was annealed with equimolar t1.5 ribozyme, BCy3P10 primer and 1.25 nmol of AUA triplet in 125 μL of water with 0.1% tween (80 ℃ 2 min, 17 ℃ 10 min). The annealed mixture was then placed on ice, 125 μL of chilled 2x extension buffer (100 mM Tris-Cl pH 8.3, 400 mM MgCl_2_) was added, and the reaction was frozen with dry ice and incubated at -7 ℃ overnight. The reactions were then stopped using equimolar EDTA to chelate magnesium ions. This was repeated using the 12×6AUA library under the same conditions.

A similar setup was used for the 6×6GAA and the 12×6GAA libraries. Key differences involve adding GAA triplet instead of the AUA triplet and incubating the reaction at -7 ℃ for 3 hours instead of overnight.

The biotinylated RNA was isolated from the reactions by binding to streptavidin coated paramagnetic beads (Dynabeads MyOne Streptavidin C1, Invitrogen). Beads were washed twice in BWBT buffer (0.2 M NaCl, 10 mM Tris·HCl pH 7.4, 1 mM EDTA, 0.1% Tween-20), twice in NaBET25 (25 mM NaOH, 1mM EDTA, 0.05% Tween 20), twice in BWBT, and then resuspended in 95% Formamide with 25 mM EDTA. This suspension was heated at 94°C for 5 minutes to elute the biotinylated products from the beads. The eluate was loaded on a denaturing PAGE gel to separate the full-length product from the primer and intermediate products. The full-length products were PAGE purified and extracted from the gel fragments onto streptavidin coated paramagnetic beads. The beads were isolated from the gel slurry by filtering of the supernatant using 50 μm size filters (Partec Celltrics, Wolflabs). The beads were then transferred to a series of reactions to enable the amplification of the bead-bound sequences. Namely, the 2′,3′-cyclic phosphate resulting from HDV cleavage was removed using T4 PNK (NEB). A constant adaptor sequence (AdeHDVlig) was then ligated to the 3′-end of the bead-bound recovered sequences using T4 RNA ligase 2 truncated KQ (NEB). The ligated product was then used as template for RT-PCR using SuperScript III One-Step RT-PCR System with Platinum Taq DNA Polymerase (Invitrogen), with primers HDVrec and forceGG at 0.5 μM each. The library prior to the selection was also PNK treated, adapter ligated and RT-PCR amplified in parallel, with no purification in between each step (5 pmoles of RNA was used in a 50 μL T4 PNK reaction, 0.1 pmoles were carried forward in an adaptor ligation reaction, and 0.01 pmoles were used for RT-PCR).

The RT-PCR products were further amplified to introduce the P7 and P5 Illumina sequencing adapters by PCR (primers used for each library are detailed in **Supplementary Table 1**) using GoTaq Hot Start (Promega). The PCR products were agarose gel purified (Qiagen), quantified using Qubit dsDNA HS Assay Kit (Invitrogen) and sequenced using a MiSeq sequencer using a 600-cycle v3 chemistry kit set-up for 2×250 reads. The reads were paired and quality filtered using fastp^59^ and demultiplexed using cutadapt^60^. The demultiplexed reads were analysed with python scripts available at github.com/chaitjo/geometric-rna-design.

### Fitness calculation

The fractional abundance of each variant in the pre-selection (input) and post-selection (output) libraries was determined by next-generation sequencing as described above. Fitness was calculated as the log2 enrichment of a variant relative to the wildtype 5TU sequence. Sequences with at least 5 reads in the pre-selection library and at least 1 read in the post-selection library were classified as active if the fitness was above an activity threshold. Sequences with zero reads in post-selection libraries which cannot be assigned a fitness value were classified as inactive.

We used the 6×6AUA libraries with a short linker incubated overnight for the results in the main text (**Figure 3**). These libraries were chosen based on obtaining the highest Pearson correlation coefficient of 0.86 between the average per-junction ligation efficiency from a low-throughput gel assay (described subsequently) and the high-throughput fitness (**Figure 3F** and **Supplementary Figure 4**). We used the high-throughput fitness of the least active variant 319 (as determined by the gel) as our activity threshold at -1.86.

### Primer extension low-throughout assay of ribozyme variants

In order to assay the activity of 5TU and its gRNAde-designed variants in detail, we individually transcribed and purified each RNA sequence (as described in Preparation of RNA constructs). The PAGE purified RNA was used to set-up primer extension reactions where the RNA variants were challenged to copy a template using triplet substrates (see **Supplementary Figure 2** for a visual depiction). The reaction was set-up by mixing primer, template, 5TU ribozyme variant, t1.5 subunit, in 0.1% Tween-20 in half the final reaction volume and heat annealing (80 ℃ 2 min, 17 ℃ 10 min). The remaining reaction volume containing 2x chilled reaction buffer (100 mM Tris-Cl pH 8.3, 400 mM MgCl_2_) was added to the reactions, which were then frozen in dry ice and incubated at -7 ℃ for the time indicated in a R4 series TC120 refrigerated cooling bath (Grant). The primer extension was separated using denaturing PAGE, and visualized on a Typhoon Trio scanner (GE Healthcare). The band intensity values were measured using ImageQuantTL (Cytiva) and used to compute the per-junction ligation efficiency, as following:

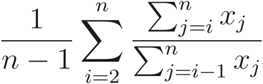

### where x_j_ is the band intensity corresponding to the j^th^ triplet incorporation

For the primer extension reactions on a template encoding 14 repeats of CGU (**Figure 3F)**, we used the top 9 gRNAde variants by fitness in the 12×6 GAA high-throughput screen. Reactions were analysed on a 20% 19:1 mono:bis acrylamide denaturing PAGE gel. Reaction conditions: 0.25 μM primer F10, 0.25 μM template tP10_14CGU, 0.25 μM each 5TU variant, 0.25 μM t1.5, 5 μM pppCGU, 200 mM MgCl_2_, 50 mM Tris-Cl pH 8.3, 0.05% Tween-20, 18 h incubation at-7C. Reactions were stopped with equimolar EDTA to the Magnesium concentration and formamide to a final concentration of 75% and denatured for 5 minutes at 94 ℃. To improve resolution, a competing strand containing an unlabelled version of the synthesized product was mixed at 10-fold excess over the template strand prior to heat denaturation and loading. A similar setup was used for the primer extension reactions on a template encoding 6 repeats of GAA (**Supplementary Figure 3**) with the top 13 gRNAde variants from the 6×6 AUA high-throughput screen.

## Data and Code Availability

The gRNAde pipeline is open-source and available at github.com/chaitjo/geometric-rna-design under a permissive MIT License (doi: 10.5281/zenodo.17934975). Pre-trained model weights are available via HuggingFace at huggingface.co/chaitjo/gRNAde (doi:10.57967/hf/7251). Training datasets, experimental data from the ribozyme assays, final designed sequence libraries, and all analysis scripts used to create figures are available via HuggingFace at huggingface.co/chaitjo/gRNAde_datasets (doi:10.57967/hf/7252). The data from the Eterna OpenKnot Benchmark is publicly available at github.com/eternagame/OpenKnotAIDesignData.

## Acknowledgements

We thank Rhiju Das and the Eterna organizers for hosting the OpenKnot Benchmark and for sharing the associated experimental data. We thank Arian Jamasb, Ramon Vinãs, Charles Harris, Alex Morehead, and Rishabh Anand for their contributions to the initial version of gRNAde. We thank Bryce Clifton, Kevin Goeij, Ganesh Agam, Maria Brunderova, and Rushikesh Dasoondi for useful discussions.

CKJ was supported by the A*STAR Singapore National Science Scholarship, Qualcomm Innovation Fellowship, and University of Cambridge Dawn HPC Pioneer Project grant. SLYK was supported by the Cambridge Trust PhD Scholarship. SVM was supported by the UKRI Centre for Doctoral Training in Application of Artificial Intelligence to the study of Environmental Risks (EP/S022961/1). This work was supported by the Medical Research Council (MRC) as part of United Kingdom Research and Innovation (also known as UK Research and Innovation (UKRI)) (MRC program grant MC_U105178804) (EG, SLYK, PH). For the purpose of open access, the MRC Laboratory of Molecular Biology has applied a CC BY public copyright license to any Author Accepted Manuscript version arising.

## Author Contributions

All authors developed the methodology. CKJ implemented the gRNAde code and generated computational designs with inputs from SVM. EG and SLYK performed the polymerase ribozyme experiments. CKJ, EG, and SLYK analyzed the experimental data. CKJ, EG, SLYK, and PH wrote the paper with inputs from all authors.

## Supplementary Figures

**Supplementary Figure 1.**
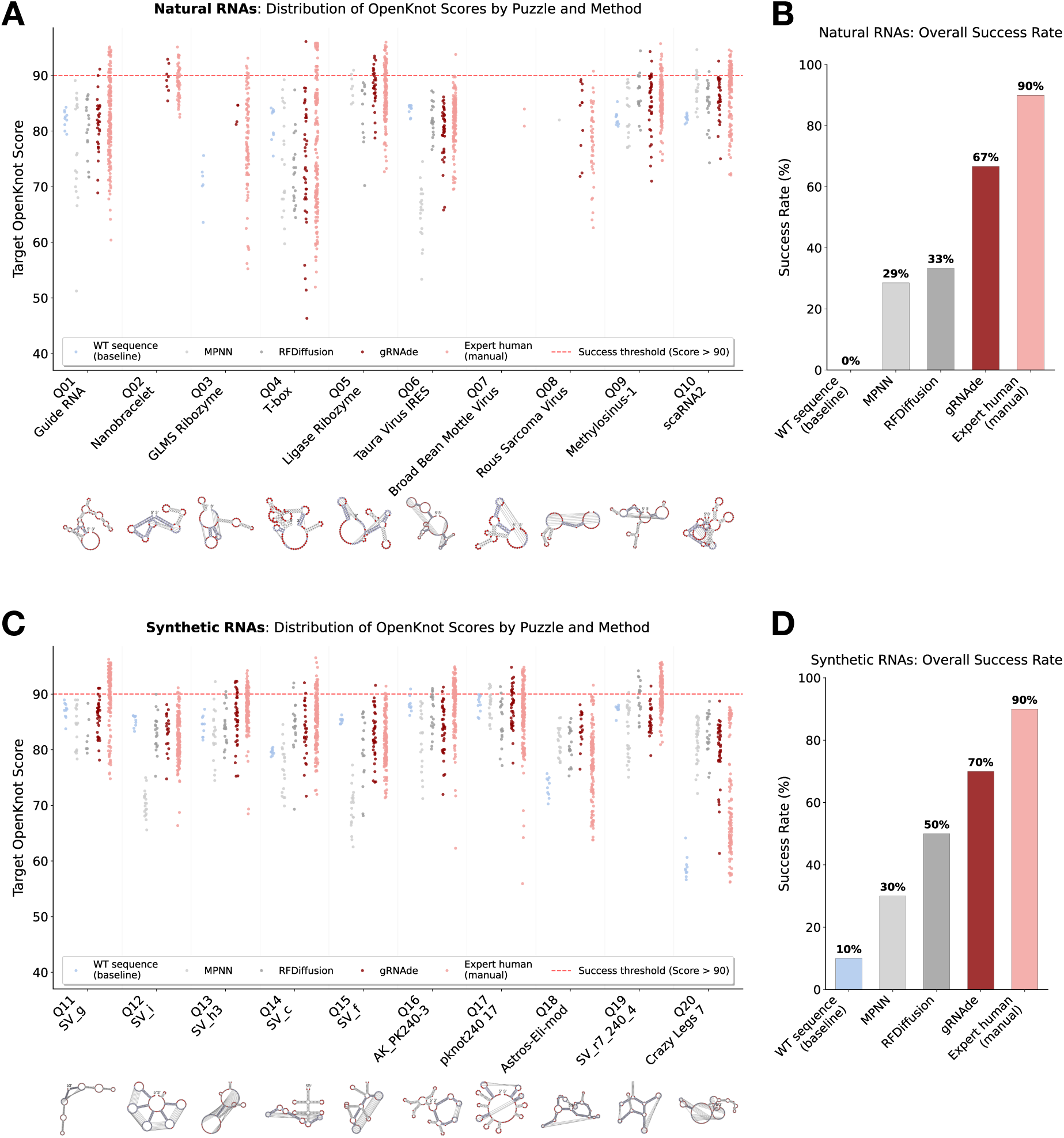
gRNAde maintains competitive performance on long RNA pseudoknots in the Eterna OpenKnot Benchmark. Performance of wildtype sequences, Rosetta^12^, MPNN^32^, RFdiffusion^31^, gRNAde, and expert human designers for the Eterna OpenKnot Round 4 challenge, which targeted pseudoknotted RNAs of up to 240 nucleotides. **A.** Distribution of OpenKnot scores for individual designs across 10 natural target puzzles (success threshold > 90, red dashed line). **B.** The success rate on natural targets, defined as the percentage of puzzles for which at least one design scored above 90. **C.** Distribution of OpenKnot scores for individual designs across 10 synthetic target puzzles (success threshold > 90, red dashed line). **D.** The success rate on synthetic targets, defined as the percentage of puzzles for which at least one design scored above 90. gRNAde achieves success rates of 67% on natural targets and 70% on synthetic targets, substantially outperforming RFdiffusion (next best AI). Rosetta (physics-based) could not be evaluated due to scalability issues. Native sequences achieved very low success rates on both categories, which demonstrates gRNAde’s ability to design idealized sequences for complex pseudoknotted RNA structures.

**Supplementary Figure 2.**
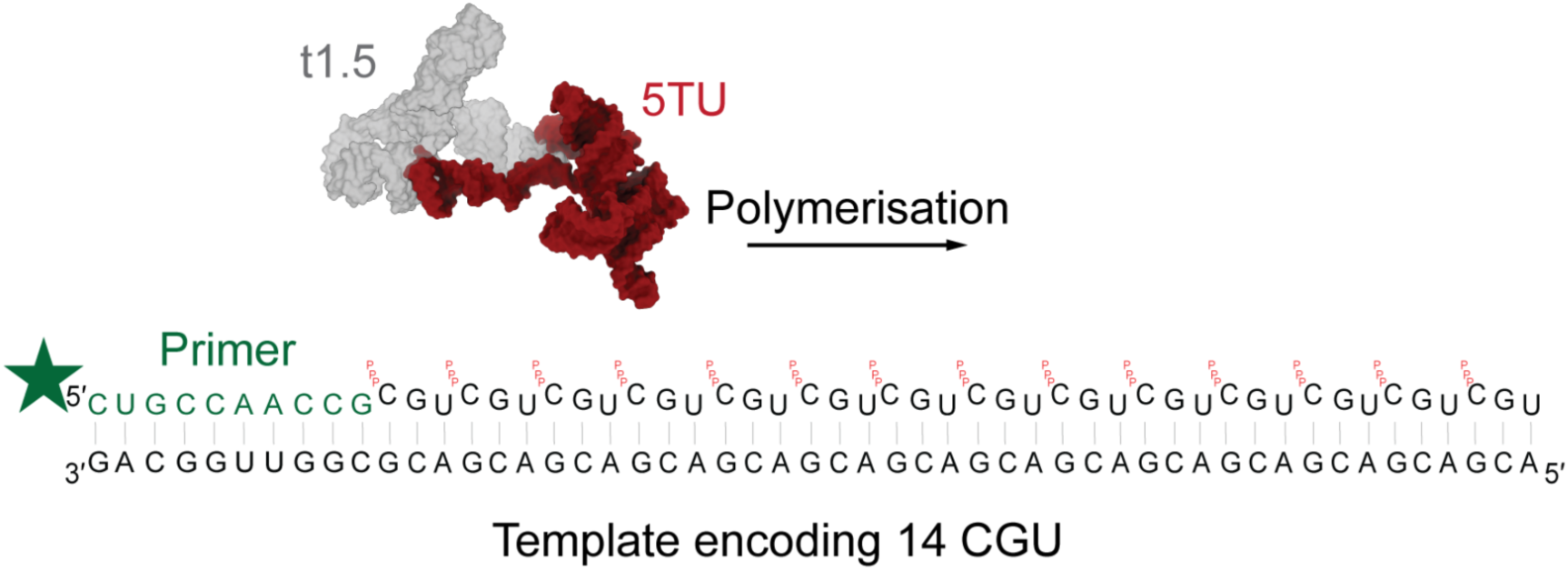
Visual representation of the primer extension process by RNA polymerase ribozymes. Synthesis of a CGU repeat template by 5TU+t1.5. A short fluorescently labelled (green star) RNA sequence primer (dark green) is extended by 5TU+t1.5 (3D representation) by one triphosphorylated trinucleotide (triplet) at the time in a template dependent manner (using template encoding 14 CGU repeats). Each triplet addition corresponds to a slower mobility band in the denaturing PAGE gel of (A), with more efficient ribozyme variants adding more triplets to the primer in the same time-frame.

**Supplementary Figure 3.**
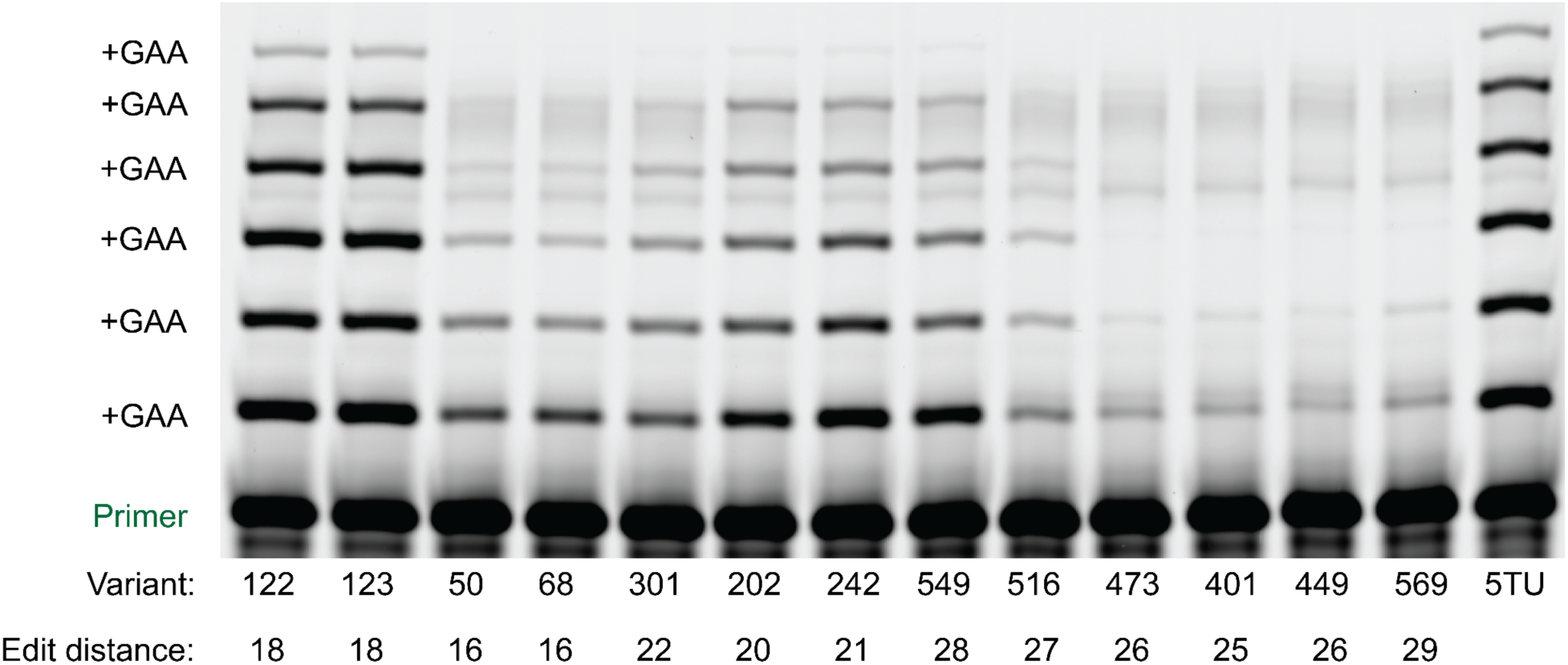
Synthesis of a GAA repeat template by gRNAde variants, showing activity at large mutational distance. Low-throughput gel analysis of primer extension reactions on a 6 GAA repeat template using the wild-type 5TU ribozyme and top 13 gRNAde-designed variants from the 6×6 AUA high-throughput screen. Variant identity and edit distance from native 5TU are labeled. The gel-confirmed activity of variant 549, which carries 28 mutations, is a key finding, proving that gRNAde can generate functional ribozymes at large mutational distances beyond those typically accessible by rational design or directed evolution. Variants 122 and 123 also show high activity, comparable to the native 5TU ribozyme.

**Supplementary Figure 4.**
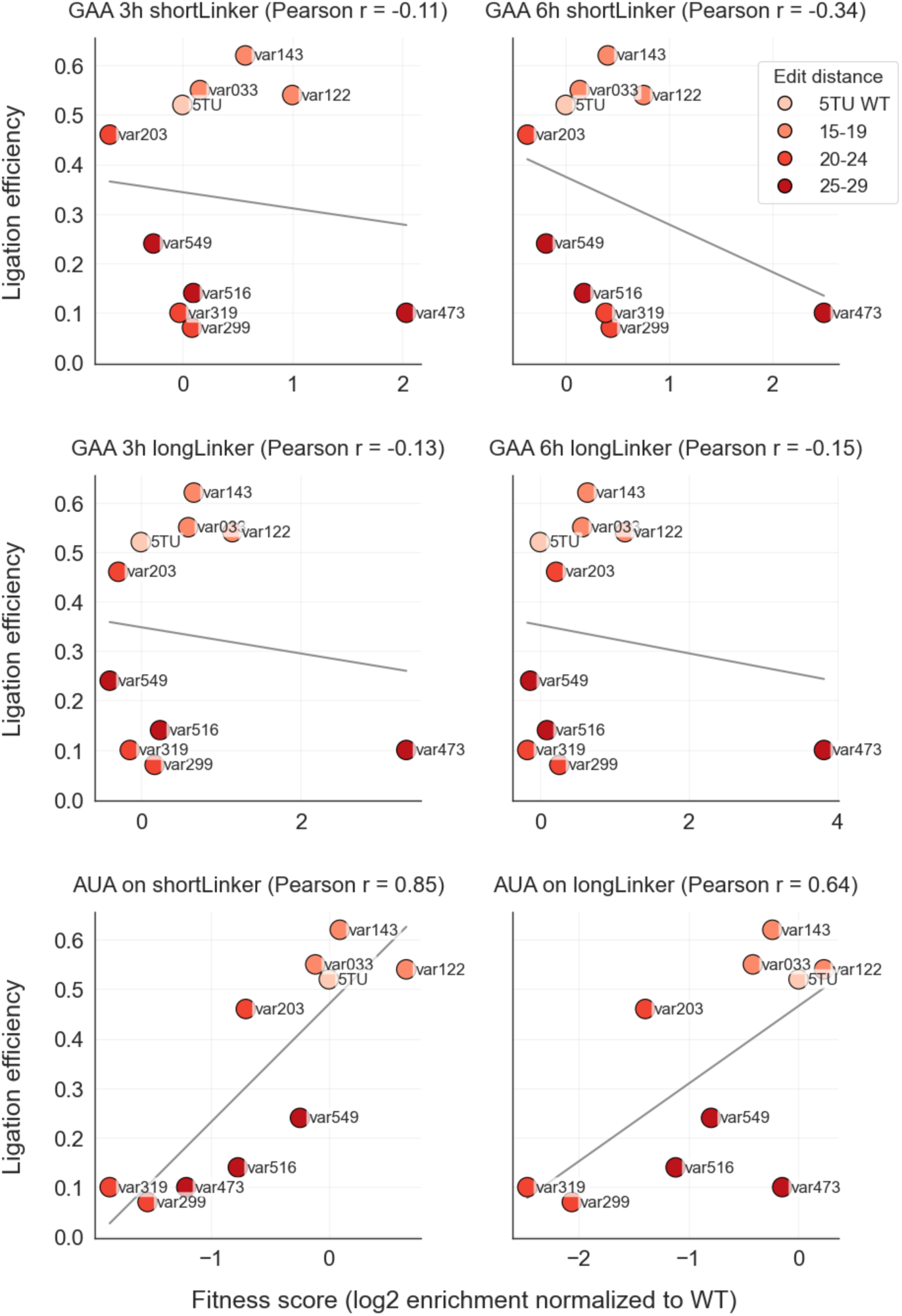
Correlation analysis of different experimental conditions for the high-throughput functional assay with the low-throughput gel assay. Scatter plots showing the correlation between the high-throughput fitness and average gel per-junction ligation efficiency (Figure 3F) for native 5TU sequence and 9 gRNAde variants under six different experimental conditions. We observe the highest correlation with the AUA libraries with a short linker incubated overnight.

**Supplementary Figure 5.**
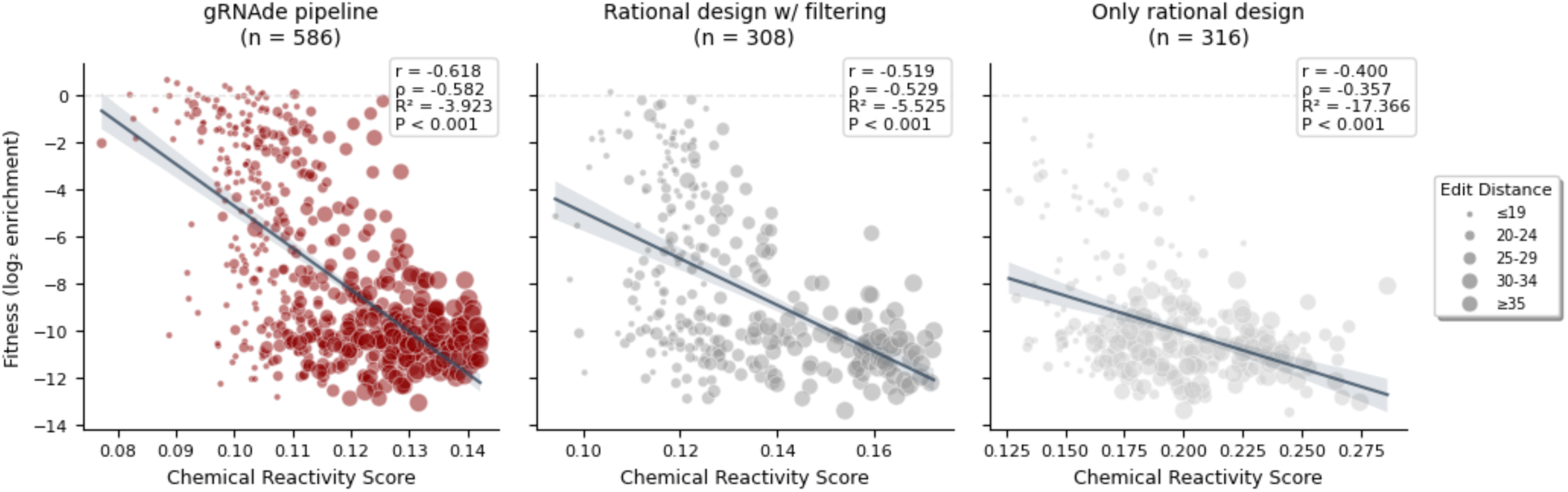
Correlation analysis of computational filtering metrics with experimental fitness. Scatter plots showing the correlation between RibonanzaNet chemical reactivity scores and experimental fitness values for designs from the gRNAde pipeline as well as rational design with and without filtering. Increasing point sizes correspond to increasing mutational distance range. We observe the highest correlation for designs from the gRNAde pipeline.

## Footnotes

1 gRNAde stands for *Geometric RNA Design*; pronounced as ‘grenade’.

